# Replaying the Tape: Comparative Genomics of Color Pattern in *Heliconius*

**DOI:** 10.64898/2026.03.23.713665

**Authors:** Christopher Lawrence, Carlos Arias, Owen McMillan, Daniel Rubenstein

## Abstract

Understanding the repeatability of evolution requires disentangling the roles of constraint, contingency, and convergence in shaping phenotypic diversity. Müllerian mimicry in *Heliconius* butterflies offers a powerful natural experiment, with co-occurring species independently evolving similar wing color patterns under shared selective regimes. Here, we integrate high-throughput image-based phenotyping, genome-wide association studies (GWAS), and comparative pan-genomics to investigate the genetic architecture underlying convergent wing pattern evolution across parallel hybrid zones in *Heliconius erato* and *H. melpomene*. Using automated computer vision pipelines, we extracted and quantified color pattern variation from over 650 butterfly specimens. Principal component analysis (PCA) of recolorized, landmark-aligned wing images captured biologically meaningful axes of variation, which were used as phenotypes in GWAS. We identified strong associations at known patterning loci—including *ivory:mir193* (previously *coretex), optix*, *WntA*, and *vvl*—as well as novel regions, including a chromosome 2 inversion in *H. erato* and a gustatory receptor gene (*Gr21a*) in *H. melpomene*. Comparative analyses using a Heliconius pan-genome revealed that while significant associations mapped to homologous regulatory regions across species, the specific variants were lineage-specific, consistent with parallel evolution via distinct cis-regulatory changes. These findings demonstrate that repeated adaptive outcomes can arise through different genetic paths within conserved regulatory architectures. More broadly, our study highlights the power of integrating machine learning, high-resolution phenotyping, and comparative genomics to dissect the molecular basis of convergent evolution in natural populations.

## Introduction

One of the most enduring questions in evolutionary biology concerns the extent to which evolution is predictable. Do similar selective pressures lead to similar evolutionary outcomes, or is the trajectory of life shaped largely by historical contingency (Gould 1992)? In contrast, others have proposed that evolution may be repeatable, with the same genes or pathways repeatedly targeted by selection across independent lineages (Stern and Orgogozo 2009). Disentangling these views requires empirical studies that examine the genetic mechanisms underlying convergent traits across repeated evolutionary scenarios.

Mimicry provides a powerful model system for testing the repeatability of evolution because it has evolved independently across diverse taxa, often in response to similar ecological pressures (Jamie 2017). In particular, Müllerian mimicry—where multiple unpalatable species converge on a shared warning signal—is driven by strong, positive frequency-dependent selection (Müller 1879; Mallet and Joron 1999). This creates a scenario in which phenotypic convergence is not only possible but adaptive, often leading to strikingly similar color patterns among distantly related species (Willmott et al. 2017; Pérochon et al. 2025). Because mimicry evolves repeatedly in geographically distinct populations and across taxonomic boundaries, it offers a valuable opportunity to test whether the same genetic solutions are consistently reused in response to similar selective forces.

Among the best-studied systems for studying mimicry is the *Heliconius* genus, a diverse group of Neo-tropical butterflies that exhibit remarkable wing pattern variation associated with Müllerian mimicry (Sheppard et al. 1997; John R. G. Turner 1971; J R G Turner 1981). Across the genus, distinct species and populations often converge on nearly identical warning color patterns, frequently in parallel across different geographic regions (Rueda-M et al. 2021; Pereira Martins et al. 2022). These patterns are not only visually striking but are also controlled by a relatively simple and modular genetic architecture (McMillan et al. 2020; Van Belleghem et al. 2023). A handful of major loci—such as *optix*, *wntA*, and *ivory:mir193 (*previously *cortex)*—control key aspects of forewing and hindwing patterning and have been repeatedly co-opted across species and populations to generate convergent phenotypes (Lewis et al. 2019; Van Belleghem et al. 2023; Livraghi et al. 2021). These loci exhibit features that facilitate their evolutionary reuse, including modular *cis*-regulatory elements that allow for fine-tuned spatial control of color patterning. However, recent findings have challenged this assumption, showing that several color patterning genes can act pleiotropically and may influence other traits such as UV reflectance and structural coloration (Mazo-Vargas et al. 2017, 2022; Thayer et al. 2020). This raises the possibility that constraints arising from shared genetic architecture may influence the range of phenotypes that can evolve.

Parallel hybrid zones within Heliconius species offer natural replicates for studying how selection, gene flow, and genetic architecture shape convergent phenotypes (Meier et al. 2021; Montejo-Kovacevich et al. 2022, 2020; Nguyen et al. 2025). Many *Heliconius* hybrid zones occur across environmental transitions where abiotic factors such as elevation or habitat structure vary alongside shifts in predator communities and local mimicry rings (Figure 1). In these environments, predator-mediated Müllerian mimicry imposes strong stabilizing selection on warning coloration (Chouteau et al. 2016; Arias et al. 2016). Despite this strong selection, many Heliconius populations exhibit imperfect mimicry, where wing patterns only approximate those of co-mimetic species (Van Belleghem et al. 2020). For example, among the four subspecies examined in this study, subtle differences occur in the hindwing rays and in the position of orange elements within the proximal forewing band (Figure 1).

**Figure 1.**
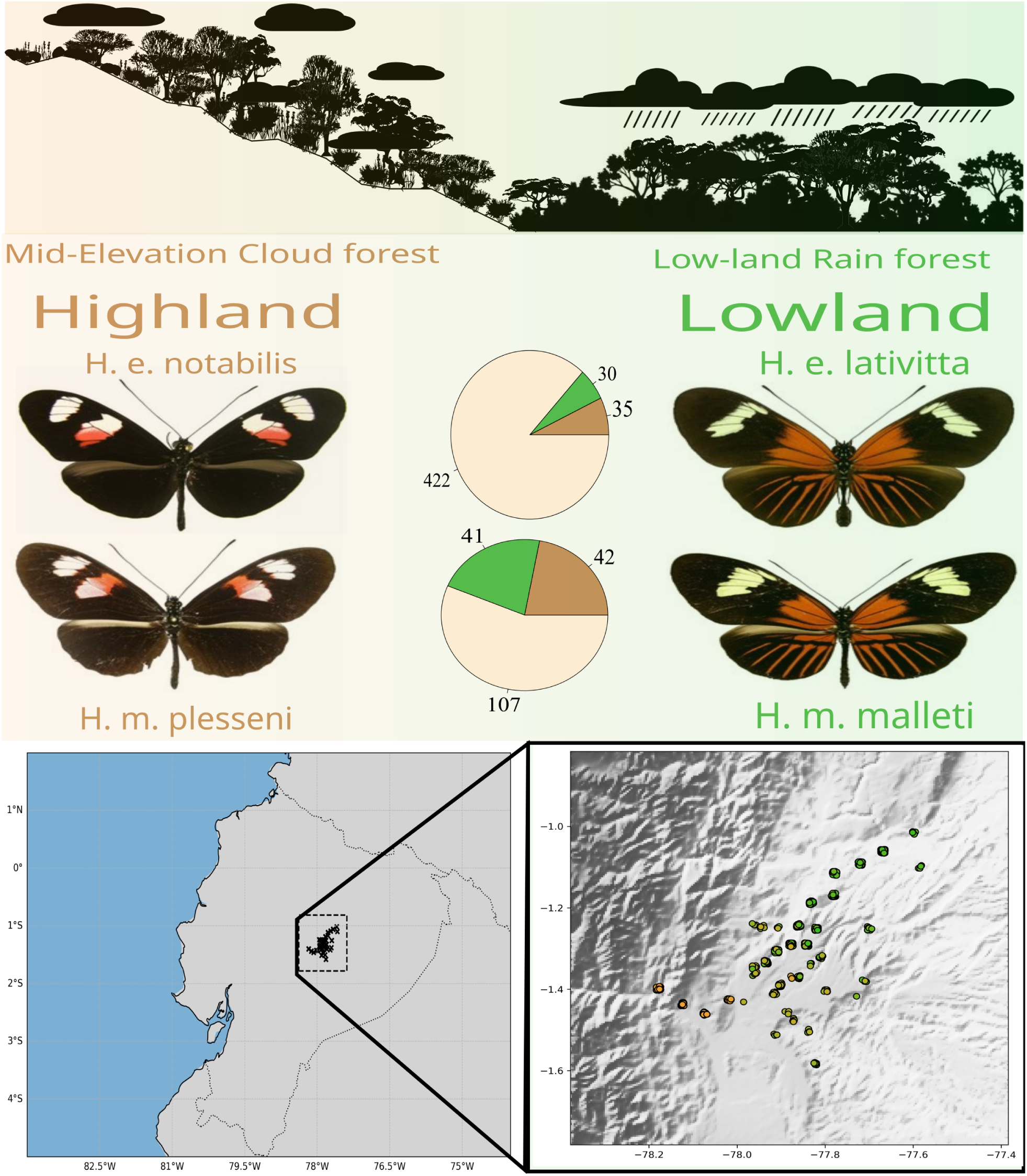
Ecological and geographic of the parallel Heliconius hybrid zones in Ecuador. This figure illustrates the ecological gradient from the mid elevation cloud forest (orange) to wet tropical rainforest (green) across which hybridization occurs between highland and lowland subspecies of *Heliconius erato* and *Heliconius melpomene* in eastern Ecuador. Pie charts represent the proportion of sampled individuals of each parental phenotype and their hybrids, with hybrids denoted in tan. The map in the bottom right shows the geographic distribution of individuals sampled in the study, colored according to phenotype status.

The persistence of such variation raises an important question: what evolutionary processes generate and maintain imperfect mimicry despite strong selection for convergence? One possibility is *de novo* mutation, which can introduce novel variation in wing pattern elements (Jiggins et al. 2017). Alternatively, introgression between hybridizing populations may transfer patterning alleles or modifier loci across species boundaries (Edelman et al. 2019; Enciso-Romero et al. 2017; Moest et al. 2020). A third explanation is that genomic constraints within the wing patterning architecture may limit the range of achievable phenotypes (Van Belleghem et al. 2020). These processes are unlikely to act independently; rather, imperfect mimicry may emerge from interactions among new mutations, introgressed genetic variation, and genomic constraints. Hybrid zones therefore provide a powerful setting to examine how these forces interact, revealing how new variation arises and spreads despite strong stabilizing selection on mimicry.

Historically, studies of wing pattern relied on qualitative classifications based on visual scoring (Joron et al. 2006; Papa et al. 2013; Meier et al. 2021). However, recent advances in high-throughput phenotyping—such as image-based trait quantification using machine learning and computer vision—now enable fine-scale, quantitative analysis of traits across thousands of individuals (van der Bijl et al. 2025; Karagic and Kratochwil 2025). At the same time, genomic technologies have advanced rapidly, allowing for high-resolution mapping of trait-associated loci using genome-wide association studies (GWAS), as well as improved structural and haplotypic resolution through graph-based pan-genomic approaches (Jandrasits et al. 2018; Meier et al. 2021; Goodwin et al. 2016). Together, these tools make it possible to connect phenotype to genotype with unprecedented precision and to test whether convergent phenotypes across hybrid zones are underpinned by shared or distinct genetic mechanisms.

In this study, we investigate the genetic basis of color pattern variation across a parallel hybrid zone in *Heliconius*, using this system to test the repeatability and predictability of evolutionary outcomes. We combine high-throughput image-based phenotyping to quantify fine-scale variation in wing patterns, genome-wide association studies (GWAS) to identify loci associated with this variation, and pan-genomic approaches to examine the structural landscape of candidate regions across populations. By comparing the genetic architecture of color pattern traits across a parallel hybrid zone, we ask whether the same genomic regions are repeatedly used in convergent adaptation. Our results provide new insights into the flexibility and constraints of adaptive evolution, offering a deeper understanding of how genetic architecture shapes evolutionary outcomes under similar selective regimes.

## Materials and Methods

### Study System

The *Heliconius* genus contains approximately 48 species, and a multitude of subspecies, distributed throughout the Neo-tropics. They can be found from southern Mexico into Central America and across much of South America. Within this radiation, *Heliconius erato* and *Heliconius melpomene* are among the most intensively studied species (Merrill et al. 2015; Davey et al. 2016; Seixas et al. 2021). Phylogenetic analyses indicate that these species diverged approximately 10–12 million years ago (Kozak et al. 2015, 2021). Both species exhibit extensive intraspecific geographic variation in wing patterning, with numerous subspecies distributed widely across the Neotropics (Rueda-M et al. 2021; Pérochon et al. 2025). Importantly, H. erato and H. melpomene frequently co-occur across diverse environments, forming parallel mimicry races across broad geographic regions (Mallet and Gilbert Jr. 1995). This co-occurrence spans multiple environmental gradients, from sea level to roughly 1,820 meters above sea level, and occurs across a variety of ecological conditions (Rueda-M et al. 2021). At transitions between ecological regions along these gradients, hybrid zones frequently form in both species, where divergent subspecies meet and interbreed (Jiggins et al. 1996; Montgomery et al. 2021; Montejo-Kovacevich et al. 2022; Rivas-Sánchez et al. 2024).

In this study, we analyze data collected from two pairs of subspecies occurring in a parallel hybrid zone in eastern Ecuador: Lowland (300m above sea level) subspecies: *Heliconius erato lativitta* and *Heliconius melpomene malleti* and Highland (1,820m above sea level) subspecies: *Heliconius erato notabilis* and *Heliconius melpomene plesseni* . These subspecies occupy distinct ecological habitats. The highland environment consists primarily of mid-elevation Andean cloud forest, whereas the lowland environment is characterized by Amazonian lowland rainforest (Figure 1).

### Sample Collection

Sampling was conducted along an 83-km elevational transect spanning this environmental transition. The transect included 35 sampling sites, from which 1,360 butterflies were collected across both species. Individuals were collected using hand nets. Wings were detached and photographed under standardized conditions using a DSLR camera equipped with a 100 mm macro lens. The resulting images and associated metadata are stored in the EarthCape database (https://earthcape.com/portfolio/heliconius/).

### Image Preprocessing

We downloaded all specimen images and performed manual verification to ensure dataset integrity. Each image contains a standardized arrangement of butterfly wings: a pair of forewings and hindwings positioned in four quadrants on a neutral background, typically a gray or green grid. This standardized layout facilitates automated detection and segmentation. Next, we implemented a custom machine-learning pipeline consisting of a fine-tuned YOLOv8 object detection model trained on butterfly specimen images (Varghese and M. 2024). The model detects elements contained in each image, including the ruler, label, color standard, and the forewings and hindwings. Following object detection, we applied Meta’s Segment Anything Model (SAM) to generate high-resolution segmentation masks for the detected components (Kirillov et al. 2023). SAM produces pixel-level masks corresponding to objects of interest in the image. Using these segmentation masks, each component is extracted from the original image. Extracted components are saved into separate directories according to their object class (e.g., left-forewing, right-hindwing, rulers, metadata, etc.). This automated extraction ensures consistent dataset organization and enables downstream analyses. Additional implementation details, parameter configurations, and usage instructions for the processing pipeline are provided in the Supplementary Materials.

With the wings segmented, we used a combination of OpenCV and the scikit-image Python library to rotate images to a standardized orientation and detect potential damage within the wing structures (Bradski 2000; Pedregosa et al. 2011; van der Walt et al. 2014). When possible, damaged right forewings or hindwings were replaced with their corresponding left-wing counterparts through horizontal reflection. This procedure maximized the number of usable samples retained in the dataset, yielding 473 *Heliconius erato* and 185 *Heliconius melpomene* specimens.

Next, we applied Contrast Limited Adaptive Histogram Equalization (CLAHE) using OpenCV. CLAHE enhances local contrast and improves edge definition while minimizing amplification of image noise (Zuiderveld 1994). This preprocessing step improves the visibility of wing venation patterns used for landmark detection and standardizes image illumination across specimens, thereby reducing variation introduced by differences in acquisition conditions (e.g., lighting environment, time of day, and photographer).

### Color Pattern Quantification

Wing venation provides evolutionarily conserved landmarks that are widely used in geometric morphometric analyses of Lepidoptera (Schwanwitsch 2009). We manually annotated 34 vein-intersection landmarks—18 on the forewing and 16 on the hindwing—for each segmented wing image using Fiji (Schindelin et al. 2012). These landmarks were used to align forewings and hindwings to subspecies-specific reference configurations, ensuring consistent spatial correspondence across samples.

To extract and reduce dimensionality in color data, we used the recolorize R package to perform color clustering in two stages: an initial k-means clustering into four colors, followed by refinement to three clusters (Weller et al. 2024). This approach standardizes color representation across individuals while minimizing noise introduced by specimen age, scale loss, or discoloration resulting from handling or preservation. Cluster similarity was evaluated using hierarchical clustering. Weighted mean RGB values were computed for each cluster, and the used to standardize color palettes across samples. Although natural variation exists in pigment intensity—particularly in red pattern elements in *Heliconius*—our objective was to quantify spatial variation in color pattern elements rather than within-color variation in pigmentation intensity (Dalbosco Dell’Aglio et al. 2017; Rossato et al. 2018). Standardizing color clusters therefore allowed consistent mapping of homologous color pattern elements across individuals.

Color-segmented images were rasterized and quantified using the patternize R package (Van Belleghem et al. 2018). Each wing was converted into a standardized pixel grid in which the presence or absence of each color was recorded across homologous spatial coordinates. This step enables direct comparison of color pattern across individuals. For each species and wing type, we conducted Principal Component Analysis (PCA) on the binarized color rasters. PCA summarizes spatial variation in color pattern elements by identifying orthogonal axes that capture the majority of variation across individuals. Separate PCAs were performed for each color as well as for total color pattern variation. The resulting PCA embeddings were exported as matrices with individuals in rows and principal components in columns, which were subsequently used as phenotypic inputs for genome-wide association analyses.

### Genomic Data and Processing

Whole-genome sequencing data was obtained from Meier et al. 2021, who sequenced 666 individuals: *H. erato* (n = 479) and *H. melpomene* (n = 187). Genomes were generated using the linked-read haplotagging protocol, which retains long-range haplotype information by barcoding long DNA fragments prior to short-read sequencing Meier et al. 2021. This approach enables the computational reconstruction of megabase-scale haplotypes. Mean sequencing coverage was 1.54× for *H. erato* and 2.77× for *H. melpomene*. SNP phasing and filtering procedures are described in Meier et al. 2021 and summarized in the Supplementary Materials of that study (Note S1). The final filtered datasets contained 25.4 million SNPs for *H. erato* (66.3 SNPs/kbp) and 23.3 million SNPs for *H. melpomene* (84.7 SNPs/kbp). We further filtered the VCF files to retain only individuals for which usable forewing and hindwing images were available.

### Genome-Wide Association and Comparative Genomic Analyses

We used *pca* and *fst* functions in Plink to calculate and control for population structure, and to calculate genome wide FST, respectively (Purcell et al. 2007). GWAS was performed using GEMMA v0.98.1, implementing a univariate linear mixed model (LMM) taking into account population structure (Zhou and Stephens 2012). To identify local enrichment of association signals, we applied a sliding window analysis across the output association files. GEMMA association statistics were aggregated into 50 kb windows with a 5 kb step size, and candidate windows were defined based on the local accumulation of low p-values.

Within each candidate region, we applied a Bonferroni correction using the genome-wide number of SNPs to define significance thresholds (α=0.05/N). SNPs exceeding the Bonferroni threshold were retained. Significant SNPs were annotated with nearby gene models from the H. erato demophoon v1 (Van Belleghem et al. 2017) and H. melpomene 2.5 (Davey et al. 2016) genome assemblies. For each region SNPS overlapping with genes were identified using GTF annotation files for each assembly.

### Pan-genome Alignment and Coordinate Translation

A pan-genome provides a unified reference framework that captures both homologous and lineage-specific sequences across multiple genomes (Brockhurst et al. 2019; Van Belleghem et al. 2023). This approach allows genomic coordinates to be compared directly between species and enables the identification of orthologous loci underlying shared phenotypes. Because our study aims to test whether similar wing pattern traits in *H. erato* and *H. melpomene* arise from reuse of homologous genetic variants or from independent mutations, we constructed a pan-genome to identify regions of conserved genomic structure between the two species.

We constructed the pan-genome following the approach described in (Van Belleghem et al. 2023). Briefly, we aligned the *Heliconius erato demophoon* v1.0 reference and the *Heliconius melpomene melpomene* v2.5 reference using seq-seq-pan (Jandrasits et al. 2018), which employs the progressive Mauve aligner to generate locally collinear blocks (LCBs) and non-homologous segments (Darling et al. 2004). The *H. e. demophoon* genome was prioritized as the primary reference to preserve coordinate order.

Lineage-specific intervals were identified by recording per-genome coordinates relative to the pan-genome using seq-seq-pan_blocks_intervals.py and intersecting these with a merged library of all sequences via BEDTools v2.27.1 (Quinlan and Hall 2010). We then assessed the overlap of significant GWAS loci between *H. erato* and *H. melpomene* by identifying SNPs and candidate genes falling within homologous genomic intervals in the pan-genome alignment using the mapping utility in seq-seq-pan. A locus was considered homologous if it mapped to the same LCB or orthologous gene in both species. Shared LCB regions containing significant SNPs in both species were subsequently analyzed and visualized using custom scripts.

## Results

### Image Pre-processing and Landmarking

The automated image-processing pipeline processed 674 raw butterfly wing images across two species. After quality filtering, 16 images were excluded due to structural damage, yielding a curated dataset of 658 high-quality wing images, each subsequently annotated with 34 homologous anatomical landmarks (18 forewing, 16 hindwing), resulting in a total of 22,372 landmark coordinates for alignment. Contrast normalization using CLAHE reduced variation in image brightness and improved the visibility of wing pattern elements across specimens. This standardized and landmark-annotated dataset formed the basis for subsequent geometric alignment and color pattern quantification.

### Color Pattern Quantification

Forewing and hindwing images were aligned using landmark-based registration to standardize orientation and size, enabling pixel-wise comparisons of homologous wing regions (Figure 2). Color clustering reduced the wing color space to three dominant classes (red, black, and yellow/white), allowing consistent quantification of pattern variation across specimens. Principal component analysis (PCA) of the aligned color pattern data revealed that the major axis of variation differed slightly between species but consistently captured forewing band morphology.

**Figure 2.**
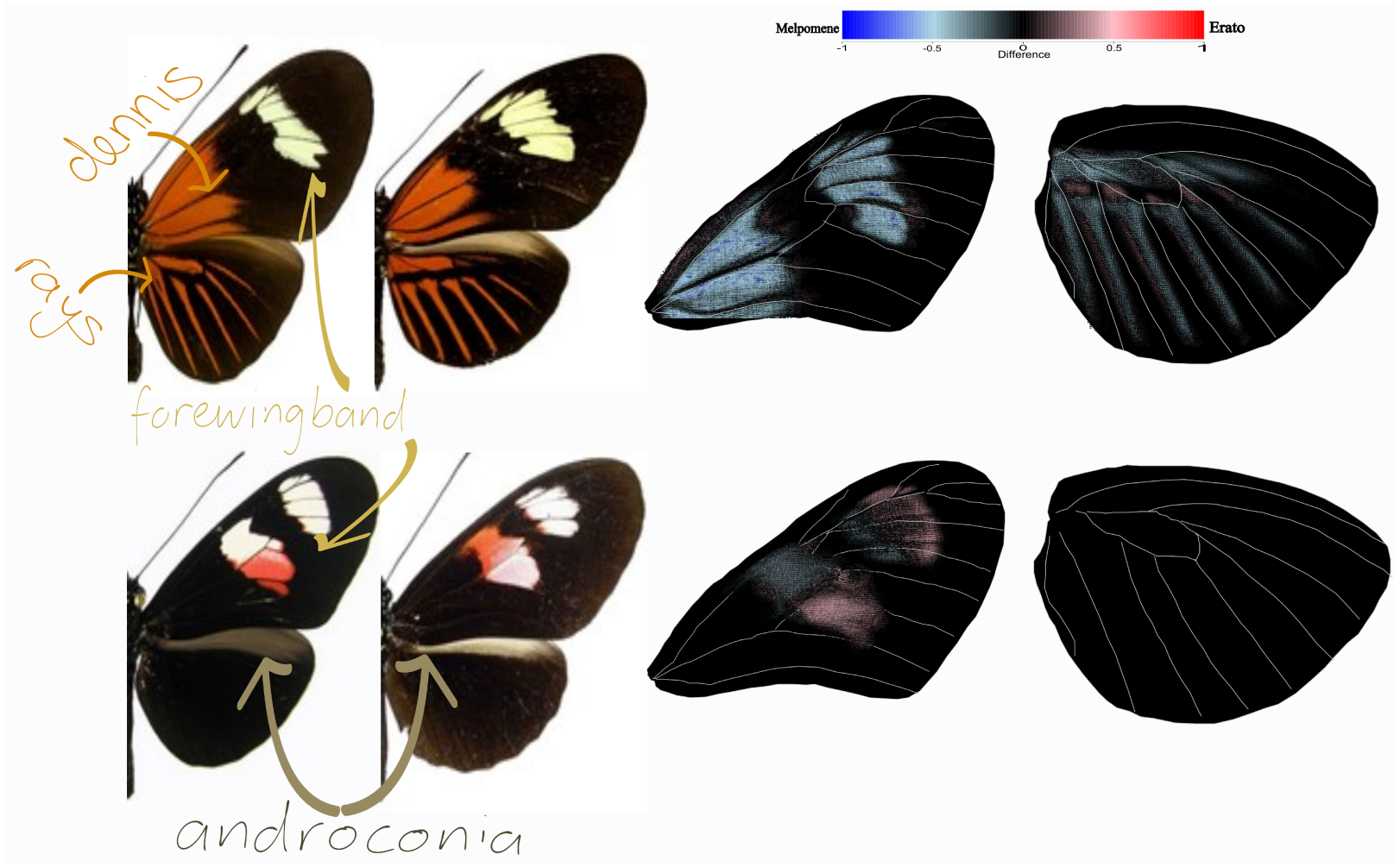
Comparative analysis of mimetic highland and lowland *Heliconius* subspecies. This figure shows color patterns of *Heliconius erato* (left) and its co-mimic *Heliconius melpomene* (right); highland (top) and lowland (bottom). Pixel-wise phenotypic comparisons between mimetic pairs visualize areas of morphological divergence in wing coloration. Warmer colors indicate pixels where *H. erato* exhibits color but *H. melpomene* does not; cooler colors indicate the reverse.

In H. *erato*, PC1 explained 23% of total variation and primarily reflected variation in the shape and position of the forewing band, whereas PC2 (10.2%) captured joint variation in the Dennis patch—a red basal forewing marking near the wing base (Figure 2)—and the distribution of red pigmentation within the forewing band. In H. *melpomene*, PC1 explained 34% of variation and represented coordinated changes in forewing band morphology and red distribution across the wing, while PC2 (10.8%) similarly described variation in the Dennis patch and forewing band redness.

Patterns of hindwing variation differed somewhat between species. In H. *erato*, PC1 explained 8.3% of total variation and primarily captured presence–absence variation in the red hindwing band, whereas PC2 (6.7%) reflected variation in grey coloration associated with the androconial patch—a region of specialized male scent scales involved in pheromone release (Figure 2), distinguishing male and female wing phenotypes. This axis of variation was more pronounced in H. *melpomene*, where PC1 explained 26.2% of variation and similarly captured presence–absence variation in the red hindwing band, while PC2 (12.7%) represented grey coloration associated with androconial patch size and sexual dimorphism. Detailed hindwing PCA results and subsequent GWAS analyses are presented in the Supplementary Materials.

Although the variance explained by individual principal components was modest, this is expected for high-dimensional pixel-based pattern data, where variation is distributed across many spatial features. Across both species, PC1 consistently captured the dominant axis of red pattern variation and was therefore retained as the primary quantitative phenotype for downstream genome-wide association analyses (Figure 3).

**Figure 3.**
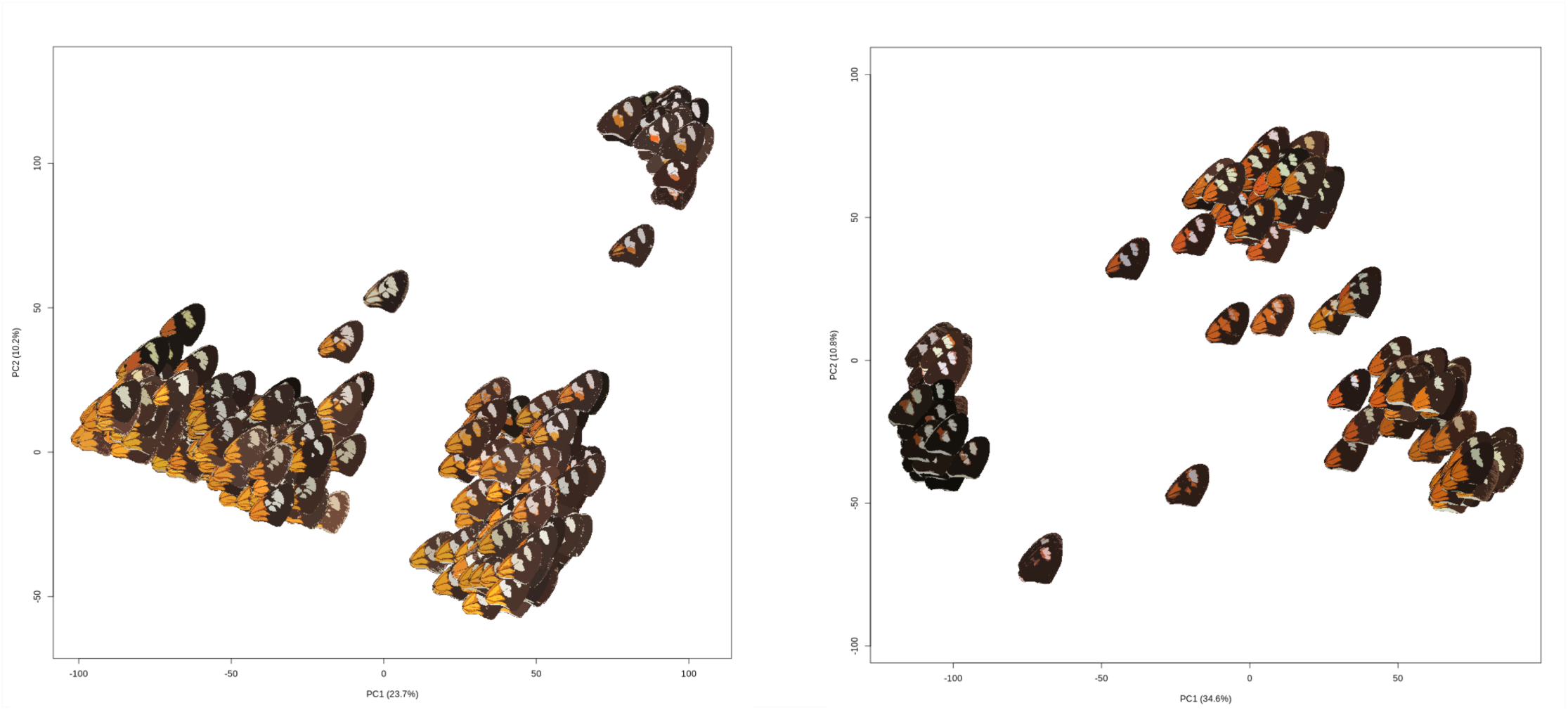
Principal component analysis of forewing color pattern variation. PCA plots summarize pixel-based variation in forewing phenotypes. **Left**: *H. erato:* PC1 explains 23.7%: PC2 (10.2%). **Right**: *H. melpomene:* PC1 explains 34.6%: PC2 (10.8%)

### Genome-Wide Association Study (GWAS)

Genome-wide association testing was performed using 49.2 million SNPs in H. *erato* and 26.3 million SNPs in H. *melpomene*. Genome-wide differentiation between highland and lowland races was low in both species (H. *erato* mean FST= 0.019; H. *melpomene* mean FST = 0.011), consistent with ongoing hybridization between neighboring populations across the altitudinal gradient. Despite this low background differentiation, association scans identified several genomic regions strongly associated with forewing color pattern variation (Figure 4).

**Figure 4.**
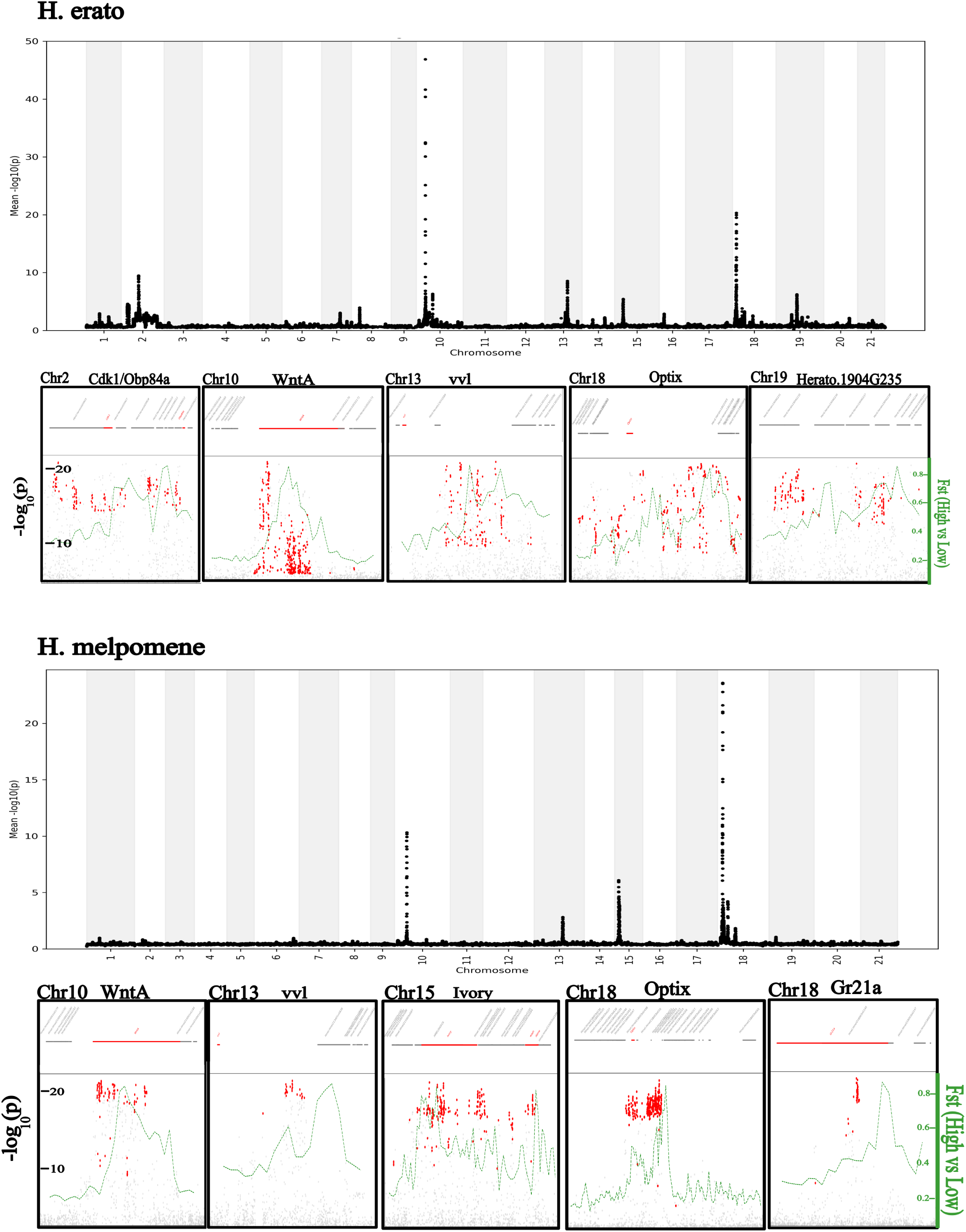
GWAS results for forewing color pattern variation in *H. erato* and *H. melpomene*. Manhattan plots show the significance of SNP associations across the genome. In both species, known color pattern loci (*optix*, *wntA*, *vvl*, and i*vory-mir193*) show strong associations. Inset sub-region plots highlight genomic intervals of interest. Significant SNPs are plotted in red; gene models are shown in grey, with previously characterized patterning genes highlighted in red. Population differentiation (fst) is overlaid as a green line.

In both species, the strongest association signals mapped to a shared set of major color pattern loci previously implicated in *Heliconius* wing pattern evolution. In H. *erato*, significant associations were concentrated in five genomic regions including three canonical patterning genes: *WntA* (chromosome 10), *vvl* (chromosome 13), and *optix* (chromosome 18). There is a peak at *ivory:mir193* on chromosome 15, but it is not significant. These loci are known to regulate forewing band shape and red pattern elements in *Heliconius*. Two additional association peaks were detected on chromosomes 2 and 19. The chromosome 2 signal overlaps a previously identified chromosomal inversion (Edelman et al. 2019; Meier et al. 2021) and includes the gene *CDK1*, which encodes a cyclin-dependent kinase involved in cell-cycle regulation and developmental processes (Jin et al. 2005). The chromosome 19 interval contains several unannotated features in the H. *erato demophoon* v1.0 assembly; however, homology searches against *Drosophila* identified a candidate gene corresponding to *CG5065*, predicted to encode an alcohol-forming long-chain fatty acyl-CoA reductase (Finet et al. 2019).

In H. melpomene, association signals similarly clustered within five genomic intervals. Four of these correspond to previously characterized *Heliconius* patterning genes: *WntA* (chromosome 10), *vvl* (chromosome 13), i*vory-mir193* (chromosome 15), and *optix* (chromosome 18). In addition, a secondary association peak was detected downstream of *optix*, corresponding to the gustatory receptor gene *Gr21a*. In *Drosophila*, *Gr21a* is expressed in CO₂-sensing neurons and contributes to behavioral avoidance responses to carbon dioxide (Suh et al. 2004; Kwon et al. 2007). These results indicate that variation in forewing color pattern is primarily associated with a conserved set of major patterning loci shared between species, but with additional species-specific signals. This pattern suggests a shared genetic framework underlying convergent wing pattern evolution, alongside lineage-specific modifiers that may contribute to species-level differences in mimetic color pattern formation.

### *Heliconius* Pan-genomics

To determine whether similar wing pattern variation in the two species is associated with the same inherited genetic variants or with different mutations occurring within the same genomic regions, we used a Heliconius pan-genome to compare three well-characterized candidate loci—*optix*, *WntA*, and *vvl*—between *H. erato* and *H. melpomene*. Specifically, we asked whether significant SNPs occur at the same genomic positions in both species (homologous sites inherited from a common ancestor) or at different positions within the same genomic regions. For each locus, we quantified the proportion of sequence shared and species-specific, identified significant SNPs located within shared locally collinear blocks (LCBs)—segments of DNA that align between species and retain the same gene order—and assessed their overlap with previously described regulatory elements (Figure 5(a-c)). The three loci showed substantial sequence divergence, with only a portion of each region shared between species. Across regions, roughly one-third of the sequence was shared, with the remainder unique to one lineage or the other (*optix*: 27.41% shared, *WntA*: 30.02%, *vvl*: 41.23%).

**Figure 5.**
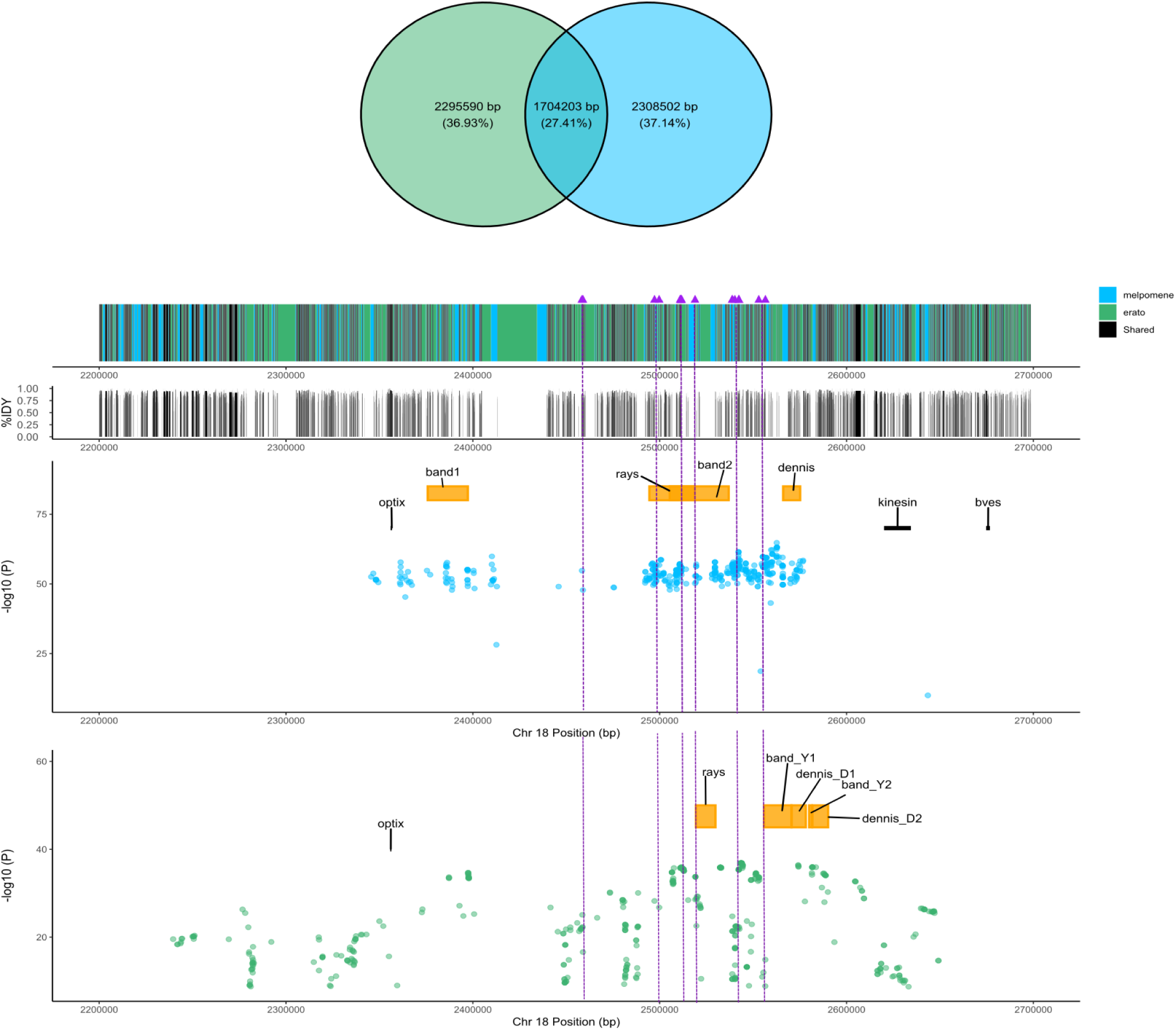
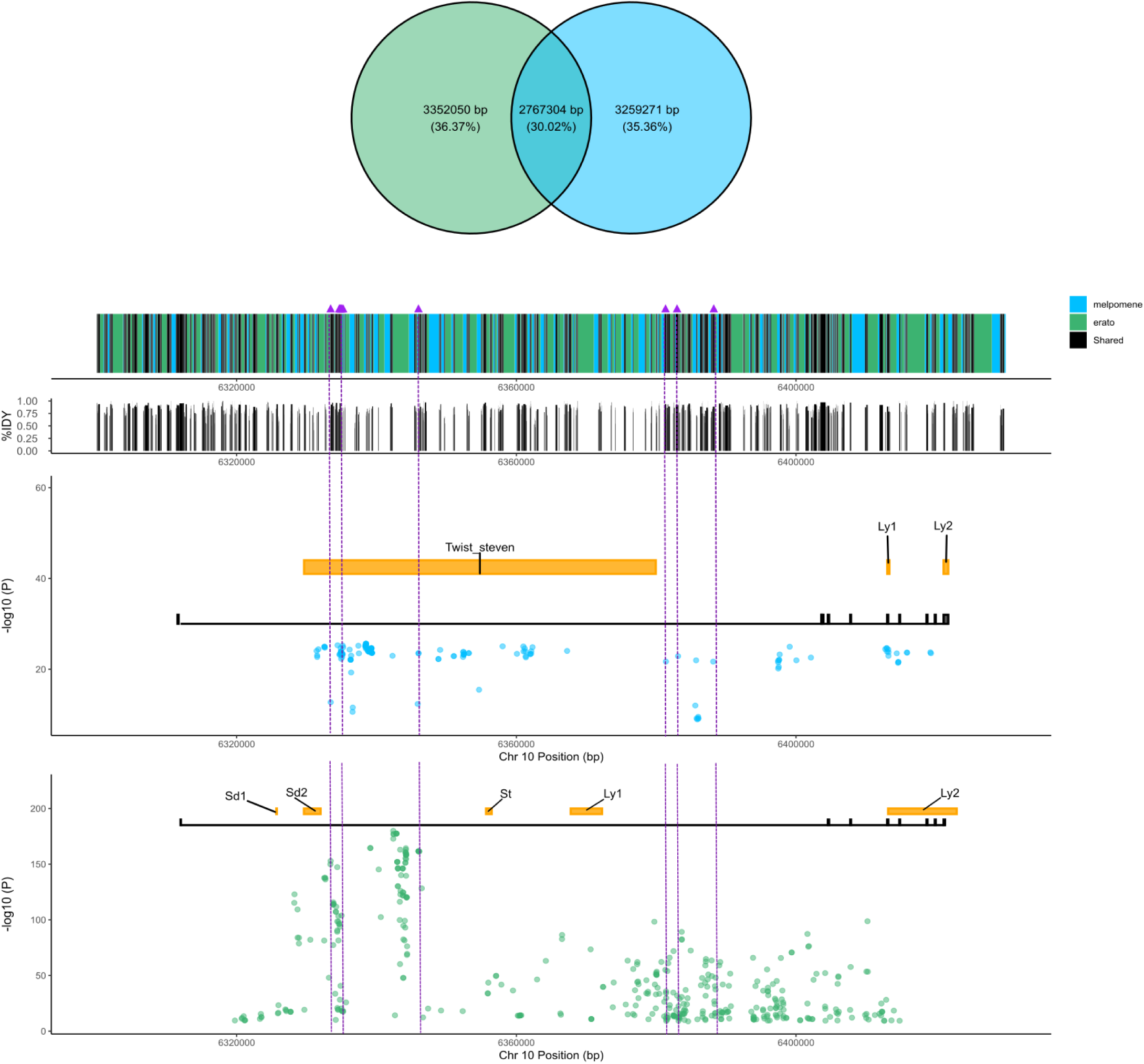
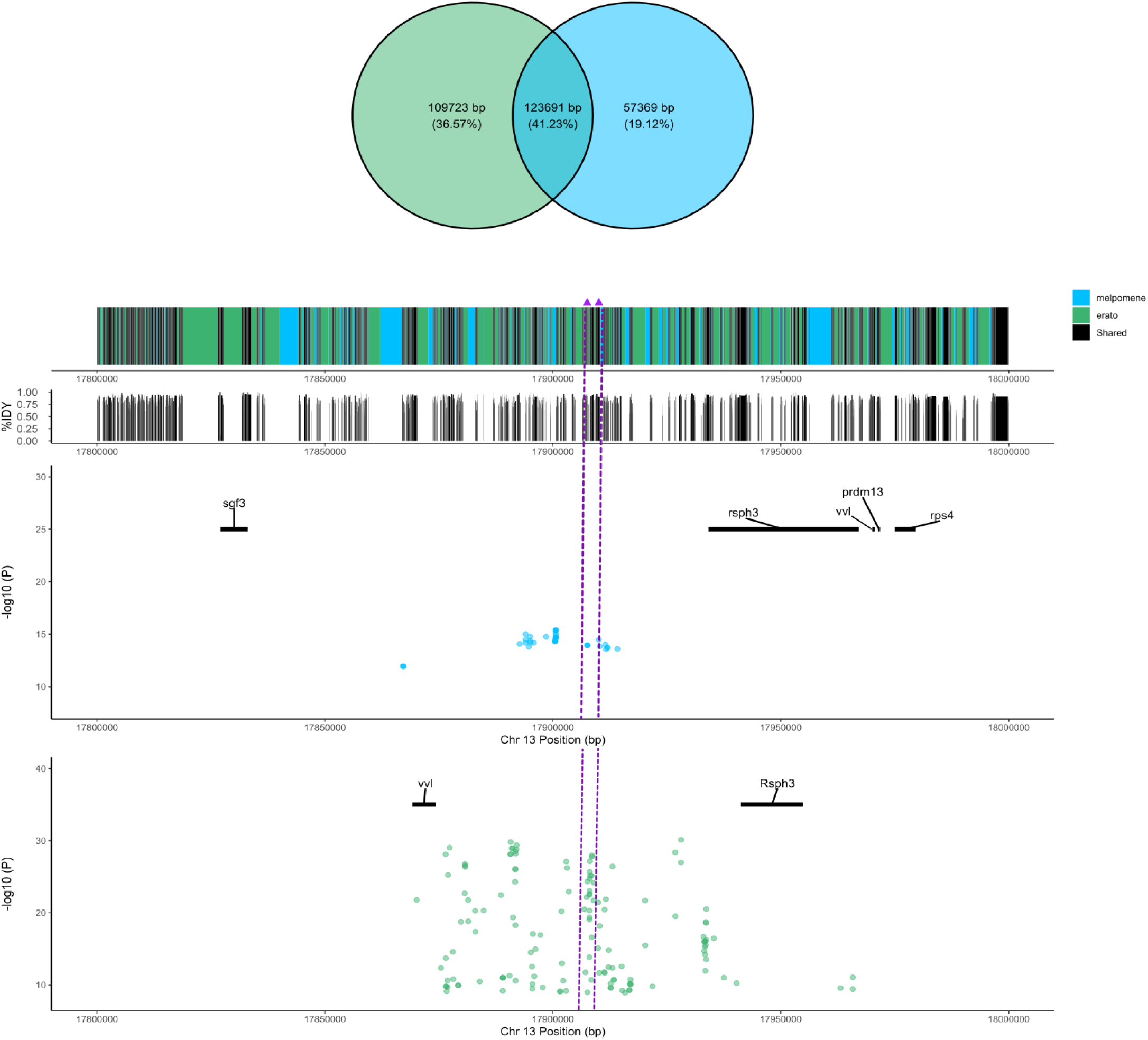
Pan-genome comparisons of major color pattern loci in Heliconius erato and H. melpomene. Pan genomic alignments of *optix*, *wntA*, and *vvl (*Figure 5*(a-c) respectively*) between *H. erato* (green) and *H. melpomene* (blue). Top row: Venn diagrams show the number and percentage of base pairs shared between species and those unique to each genome. Purple triangles denote SNPs located within shared genomic blocks. Bottom rows: Sub-region SNP plots show the distribution of associations in each species, colored by species, alongside annotated gene features. Yellow bars indicate regions previously associated with distinct color pattern elements.

At the *optix* locus, numerous significant SNPs in both species occurred within shared LCBs, distributed across 19 aligned blocks. However, no significant SNPs occurred at homologous genomic positions between species. Many of these blocks occurred upstream of *optix*, overlapping known modular regulatory elements associated with wing pattern features such as rays, band, and Dennis (Figure 5a).

The *WntA* locus showed a similar pattern. Significant SNPs were distributed across 11 shared LCBs, again with no significant SNPs occurring at homologous genomic positions between species. Several of these blocks overlapped the first intron of *WntA*, a region known to contain multiple cis-regulatory elements associated with pattern elements *SD2* and *ST*, although the spatial correspondence between SNPs and regulatory features was less pronounced than at *optix* (Figure 5b).

At the *vvl* locus, the proportion of shared sequence was highest among the three regions examined (41.23% shared). Despite this, only two shared LCBs contained significant SNPs in either species. As in the other loci, no significant SNPs occurred at homologous genomic positions between species. Structural differences were also evident across this region, with gene order rearranged in *H. melpomene* relative to *H. erato* (Figure 5c).

Across all three loci, association signals occurred within aligned genomic regions shared between species, yet the associated SNPs differed between species. Together, these results indicate that selection repeatedly targets conserved genomic loci across species, while the underlying genetic variants differ between lineages.

## Discussion

The persistence of imperfect mimicry in *Heliconius* raises an evolutionary question: do convergent phenotypes arise through shared genetic architecture or through independent genetic changes? Mimetic hybrid zones provide a powerful framework for addressing this question because phenotypically similar populations can be compared across divergent genomic backgrounds. Our results show that convergent evolution repeatedly targets the same major genomic regions in *H. erato* and *H. melpomene*, although the specific loci and mutations associated with color pattern are not identical between species. These findings suggest that convergent mimicry can arise through independent genetic changes occurring within shared developmental loci.

Empirical studies of Heliconius butterflies have repeatedly shown that convergent color pattern phenotypes are frequently produced through cis-regulatory modifications affecting a small number of major patterning genes. For example, regulatory variation around *optix* influences red pattern elements (Wallbank et al. 2016; Van Belleghem et al. 2017; Moest et al. 2020), while changes near i*vory-mir193* contribute to variation in melanic patterning (Hanly et al. 2022; Livraghi et al. 2021, 2024). These regulatory elements can act modularly, modifying discrete components of the wing pattern and facilitating the evolution of diverse mimetic forms (Enciso-Romero et al. 2017; McMillan et al. 2020; Van Belleghem et al. 2021). However, this modular view is not universally supported. Evidence from analyses of *WntA* and other patterning loci suggests that some regulatory regions exert pleiotropic effects, influencing multiple pattern elements across the wing surface (Morris et al. 2019; Lewis et al. 2019; Mazo-Vargas et al. 2017, 2022). Together, these findings suggest that the genetic architecture underlying mimetic patterns may involve a combination of modular and pleiotropic regulatory mechanisms.

Our pan-genomic comparisons provide additional context for this debate. We identified homologous sequence blocks surrounding *optix*, *WntA*, and *vvl* shared between *H. erato* and *H. melpomene*, indicating that these genomic regions were inherited from a common ancestor prior to their divergence, approximately 12 million years ago (Edelman et al. 2019; Kozak et al. 2021). Within both *optix* and *WntA*, significant associations occur in regions overlapping previously characterized cis-regulatory elements controlling color pattern. These results are consistent with a model in which similar phenotypes evolve through independent regulatory changes within conserved genomic neighborhoods, supporting parallel evolution via distinct cis-regulatory modifications (Van Belleghem et al. 2023).

An important question remains: how does cis-regulatory variation itself evolve within these genomic regions? More specifically, to what extent does color pattern evolution rely on the emergence of de novo regulatory elements versus the modification or co-option of pre-existing cis-regulatory elements (CREs)? Work by Concha et al. 2019 suggests that regulatory rewiring—the gain, loss, and reorganization of regulatory interactions—plays a central role in generating pattern divergence in Heliconius butterflies. Although pan-genomic comparisons revealed conserved locally colinear blocks surrounding major patterning genes, the specific SNPs associated with color pattern variation within these regions are not homologous between *H. erato* and *H. melpomene*. The associated variants differ in both genomic position and nucleotide identity, indicating that the causal mutations underlying pattern variation have evolved independently in each lineage. This suggests that while the broader regulatory architecture of these loci is conserved, the molecular variants generating phenotypic differences have diverged.

Additional complexity occurs at the *vvl* locus, where structural differences between *H. erato* and *H. melpomene* complicate direct comparisons of regulatory architecture. In *H. erato*, *vvl* lies upstream of *rsph3*, whereas in *H. melpomene* it occurs downstream of *rsph3* and is flanked by *pdrm13* and *rsph4*. Despite these rearrangements, both genomes retain homologous blocks containing significant—though non-homologous—SNPs associated with forewing pattern variation. This pattern suggests that even in structurally dynamic regions, selection may repeatedly target conserved regulatory neighborhoods while the specific causal variants arise independently within each lineage.

Our findings suggest that color pattern evolution involves both conserved regulatory substrates and lineage-specific molecular changes. Shared regulatory landscapes appear to channel evolutionary change toward a limited set of loci, while the mutations generating phenotypic variation arise independently within these regions. This pattern persists despite genomic divergence between *H. erato* and *H. melpomene*, including differences in genome size, structural organization, and local synteny (Cicconardi et al. 2023; Seixas et al. 2021). Such dynamics may explain why co-mimetic species converge on similar warning signals while retaining subtle but consistent differences in their wing patterns (Van Belleghem et al. 2020).

In addition to associations at well-characterized patterning loci, we also detect signals elsewhere in the genome. In *H. erato*, a significant association occurs within a previously described inversion on chromosome 2 (Edelman et al. 2019; Meier et al. 2021). Earlier work identified a QTL in this region explaining approximately 32.6% of color pattern variation, suggesting that this genomic interval may contribute to pattern diversification (Papa et al. 2013). Our analysis highlights *Cdk1*, a regulator of cell cycle progression (Jin et al. 2005), as a candidate gene within this region. Although not previously implicated directly in wing patterning, this observation is notable given parallels with the *cortex* locus, where a cell-cycle regulator influences scale development through nearby regulatory elements (Livraghi et al. 2024; Fandino et al. 2024; Orteu et al. 2024).

Beyond divergence within mimicry loci, additional ecological factors may shape genomic differentiation across the hybrid zone. In *H. erato*, we detected a second association on chromosome 19 near a gene annotated as *Dmel\CG5065*. Chromosome 19 has previously been implicated in regulatory evolution (Lewis and Reed 2019), and recent work by suggests that this chromosome contains an inversion associated with environmental adaptation, including variation in desiccation or temperature tolerance (Rueda-M et al. 2024). Because the hybrid zone examined here spans an altitudinal gradient, this signal may reflect local environmental adaptation rather than direct involvement in wing pattern determination.

A related interpretation arises in *H. melpomene*, where we detected an additional peak downstream of *optix* overlapping the gene *Gr21a*, a gustatory receptor involved in olfactory and chemosensory behavior (Suh et al. 2004; Kwon et al. 2007). Although the proximity of this signal to *optix* raises the possibility of linkage with nearby regulatory elements controlling color pattern, *Gr21a* itself is associated with sensory perception and behavior. Similar interactions between color pattern and behavior have been described near the *optix* locus: (Rossi et al. 2020, 2024) identified regions upstream of *optix*, whose neural expression correlates with male attraction to red wing patterns and whose disruption impairs conspecific courtship. These patterns highlight an important challenge in interpreting association signals in hybrid zones: loci detected in genomic scans may reflect selection acting on color pattern itself, on linked behavioral traits, or on environmentally relevant variation. Disentangling these alternatives will require additional functional and population-genetic analyses.

Although PCA can oversimplify phenotypes, the variation captured here was sufficient to identify both known and novel genomic loci (Takefuji 2025). PC1 captured 23% of the variation in *H. erato* and 34% in *H. melpomene*. The greater proportion of variation observed in *H. melpomene* is somewhat unexpected, particularly given that the dataset contains more *H. erato* specimens. One possible explanation is differences in the sampling of sexes between the two species, as both sex and geography are known to influence forewing band shape (Dalbosco Dell’Aglio et al. 2017; Rossato et al. 2018). We also conducted PCA on each color component separately; however, the resulting association signals were broadly consistent with those obtained from the combined dataset (see Supplemental Fig. XXX).

Our automated image-processing pipeline leverages recent advances in computer vision to enable high-throughput and reproducible extraction of butterfly wing phenotypes. Our segmentation model generalized well across other *Nymphalidae* species and even to more distantly related taxa such as *Papilionidae*. However, not all components of the workflow performed equally well. Automated landmark detection proved inconsistent across subspecies, particularly where wing shape and color pattern diverged substantially. This limitation likely reflects overfitting and insufficient model generalization rather than a lack of conserved anatomical reference points. This highlights both the promise and the current limitations of machine-learning–based phenotyping approaches. As image-based datasets continue to expand, analytical frameworks that integrate flexible phenotyping pipelines will be essential for scalable trait analysis (Lürig 2022).

Imperfect mimicry in *Heliconius* highlights both the predictability and flexibility of adaptive evolution. Our results show that convergent warning patterns repeatedly evolve through the same genomic regions across species, suggesting that shared developmental architecture constrains where adaptive change can occur. Yet the specific variants underlying these phenotypes differ between *H. erato* and *H. melpomene*, indicating that similar patterns arise through independent mutations rather than shared ancestral alleles. This combination of conserved regulatory landscapes and lineage-specific molecular changes helps explain how co-mimetic species achieve broadly similar warning signals while retaining subtle differences in pattern. In this sense, replaying the “tape” of evolution may produce similar phenotypes, even when the underlying genetic paths differ. As large-scale image datasets continue to expand, integrating machine learning–based phenotyping with comparative genomics will create new opportunities to test how predictable evolutionary outcomes emerge from distinct genetic routes.

