## Supplemental_materials for "Replaying the Tape: Comparative Genomics of Color Pattern in *Heliconius*"

This PDF file includes:

Supplementary Figure 1. Validation of the color quantification pipeline.
Supplementary Figure 2. PCA of hind-wing color pattern variation in *H. erato* and *H. melpomene.*
Supplementary Figure 3. GWAS of fore-wing color pattern variation in *Heliconius erato*.
Supplementary Figure 4. GWAS of fore-wing color pattern variation in *Heliconius melpomene*.
Supplementary Figure 5. Manhattan plots for total hind-wing color pattern variation.
Supplementary Figure 6. Whole-genome FST landscapes of *Heliconius erato* and *H. melpomene*.

Other Supplemental Material for this manuscript can be found here:

Lawrence, C. (2026) https://drive.google.com/drive/folders/1P90R9gErlY-OGKlEwzS7OLxjigwFU72Z?usp=drive_link

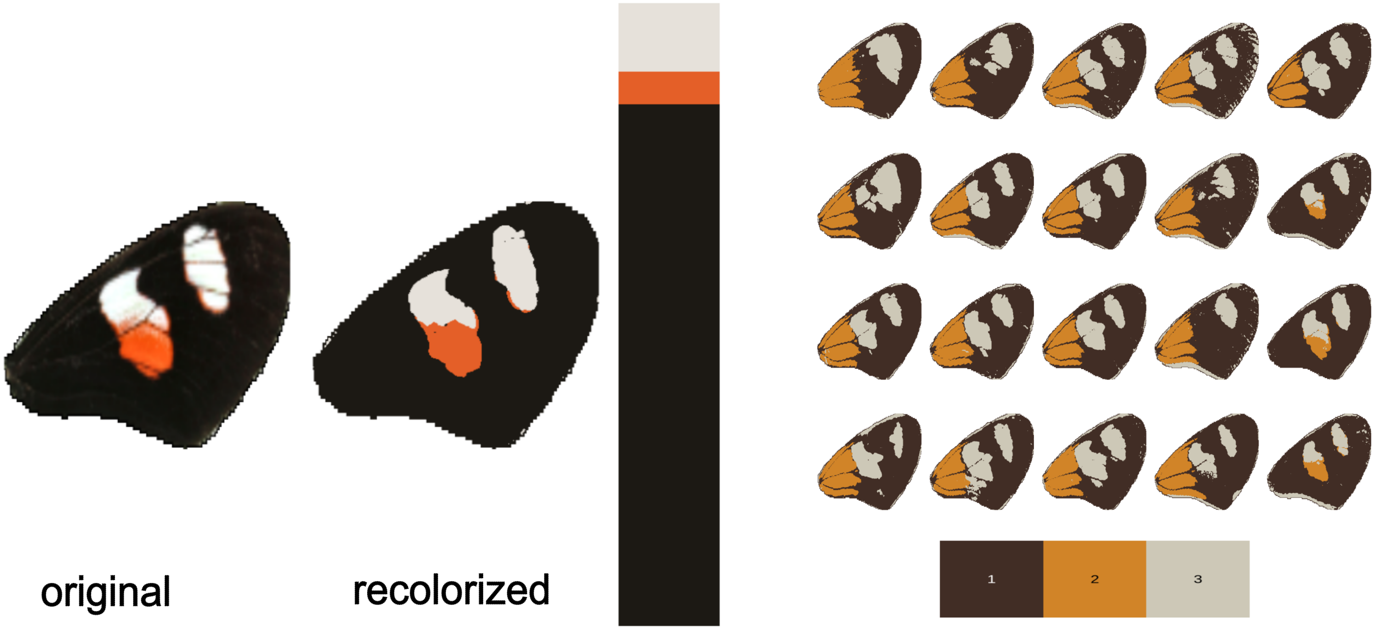


**Supplementary Figure 1. Validation of the color quantification pipeline.**
The figure illustrates the workflow used to quantify Heliconius color pattern. The original image is shown alongside the color-clustered version and the resulting isolated color clusters. A sample set of wings is then shown after mapping to the standardized color palette, with the corresponding palette displayed at the bottom. Together, these panels demonstrate the method captures and quantifies biologically relevant color pattern variation across samples.

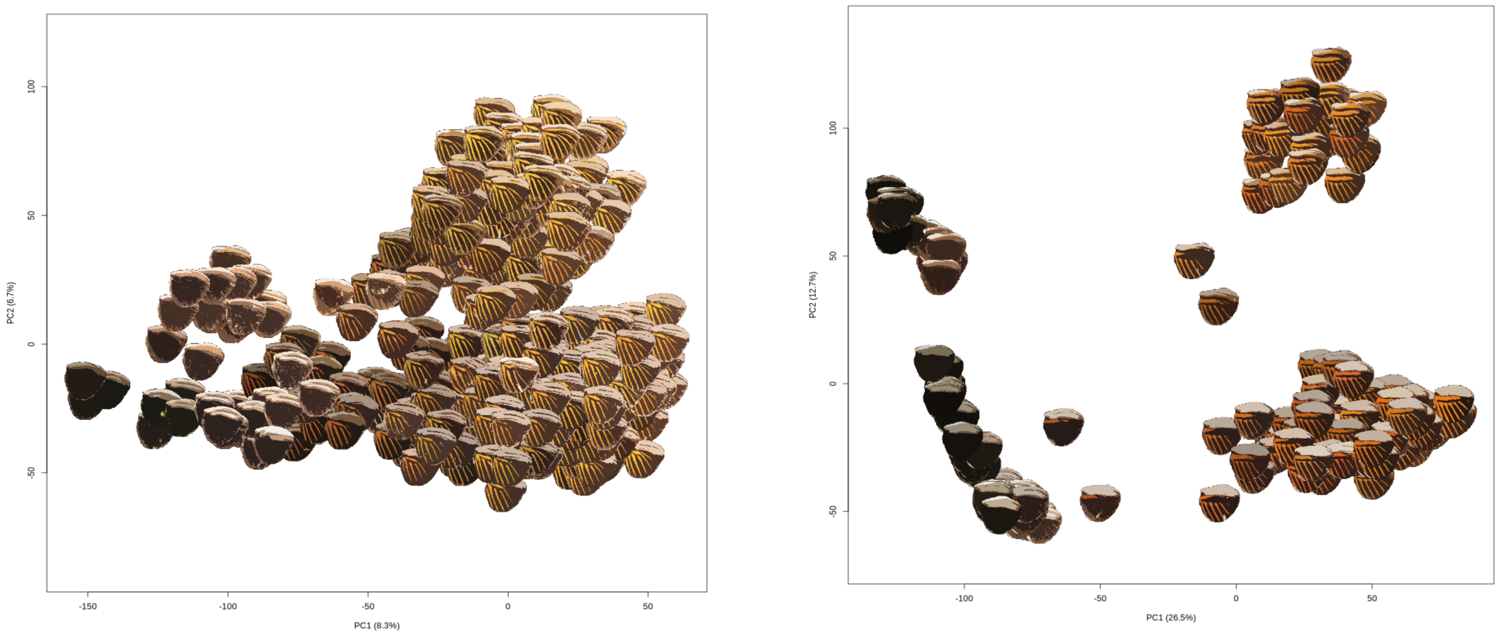


**Supplementary Figure 2. Principal component analysis of hindwing color pattern variation in *H. erato* and *H. melpomene*.**
In *H. erato*, PC1 explains 6.7% of the variation and PC2 explains 8.3% of the variation. In *H. melpomene*, PC1 explains 12.7% of the variation and PC2 explains 26.5% of the variation. In both species, PC1 primarily captures variation in the presence or absence of red "rays" on the hindwing, whereas PC2 reflects variation in the size and presence of androconial scales and captures male-female dimorphism.

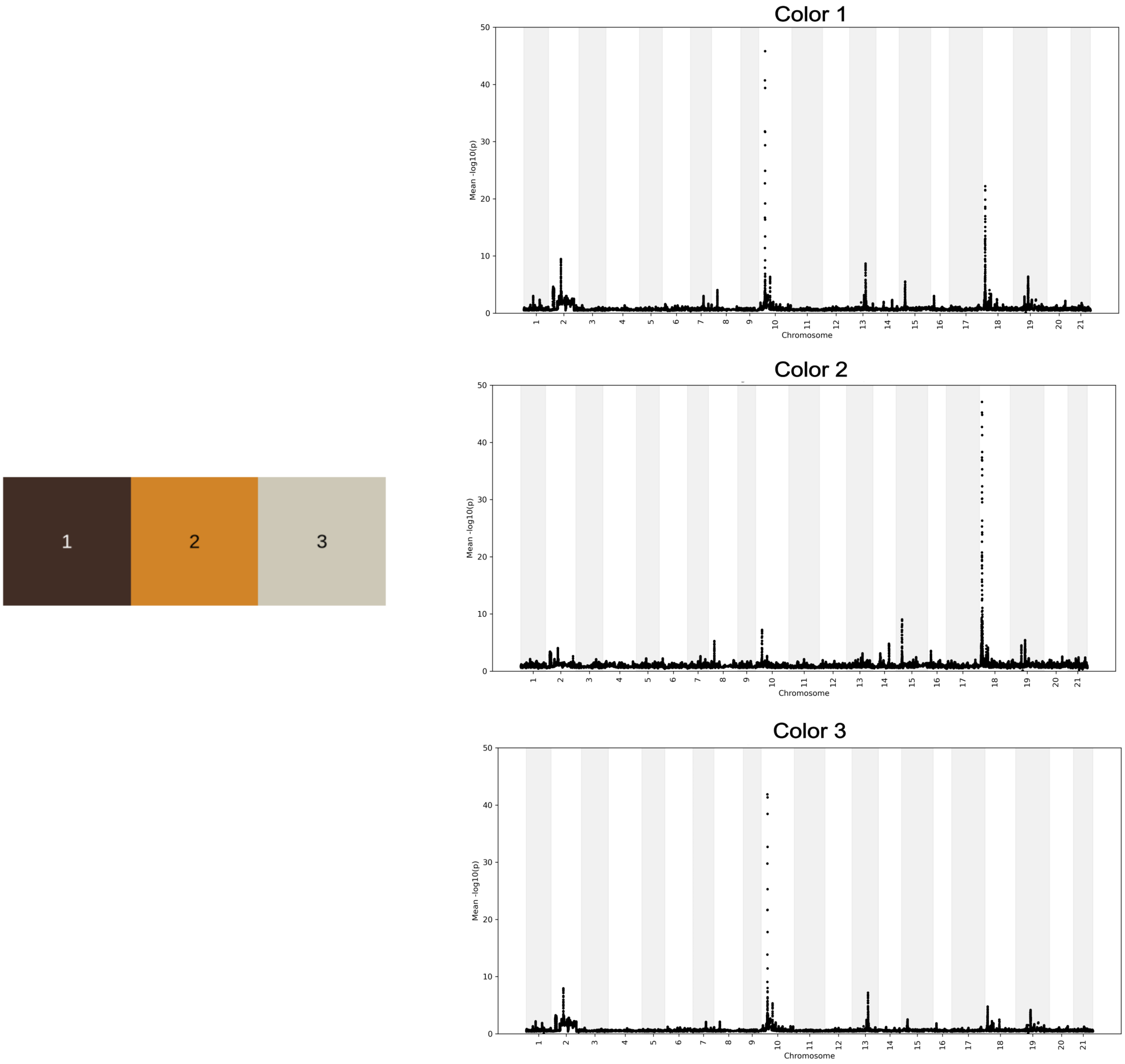


**Supplementary Figure 3. GWAS of forewing color pattern variation in *Heliconius erato*.**
The panel on the left shows the color palette used to define the focal colors. To the right, three Manhattan plots show genome-wide association results for each color separately, mapping variation in each color pattern across the *H. erato* genome. These plots identify genomic regions associated with color-specific wing pattern variation.

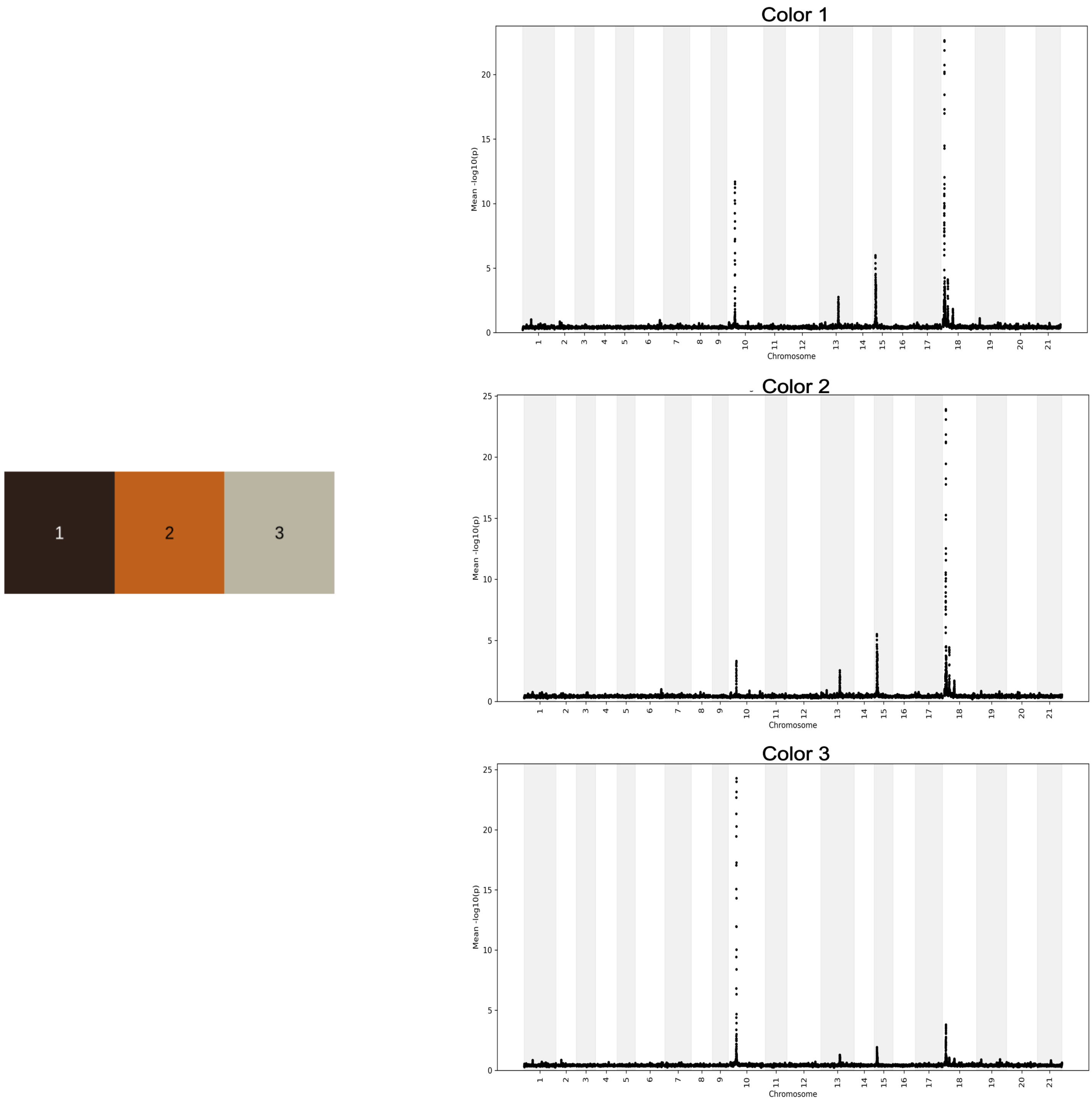


**Supplementary Figure 4. GWAS of forewing color pattern variation in *Heliconius melpomene*.** The panel on the left shows the color palette used to define the focal colors. To the right, three Manhattan plots show genome-wide association results for each color separately, mapping variation in each color pattern across the *H. erato* genome. These plots identify genomic regions associated with color-specific wing pattern variation.

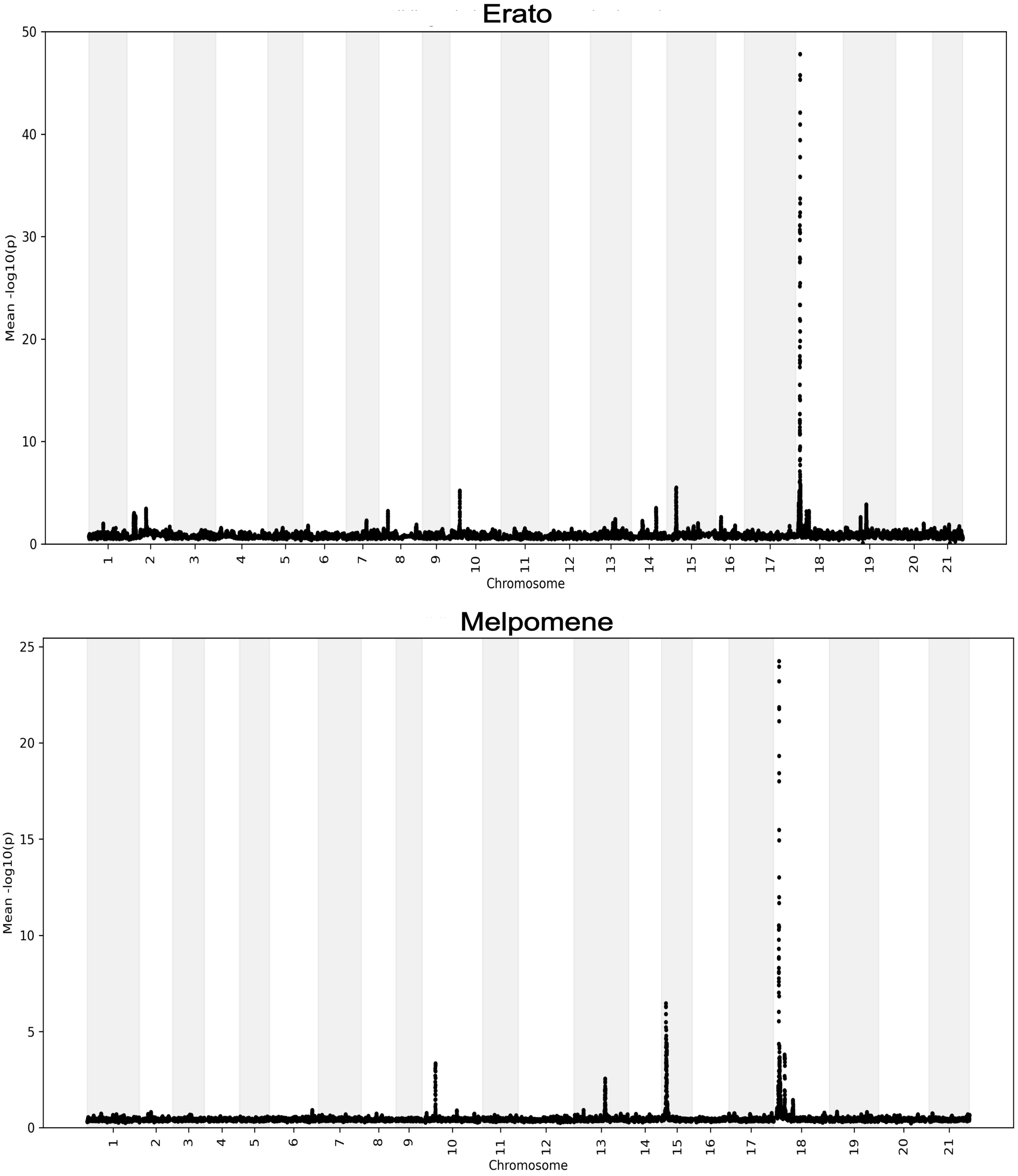


**Supplementary Figure 5. Manhattan plots for total hindwing color pattern variation in *Heliconius erato* and *H. melpomene*.**

Manhattan plots showing genome-wide mapping results for total hindwing color pattern variation in *H. erato* (top) and *H. melpomene* (bottom). The plots indicate that some of the same genomic regions and candidate genes known to control forewing color pattern are also associated with variation in hindwing patterning, suggesting partial genetic overlap in the regulation of wing pattern elements across wing surfaces.

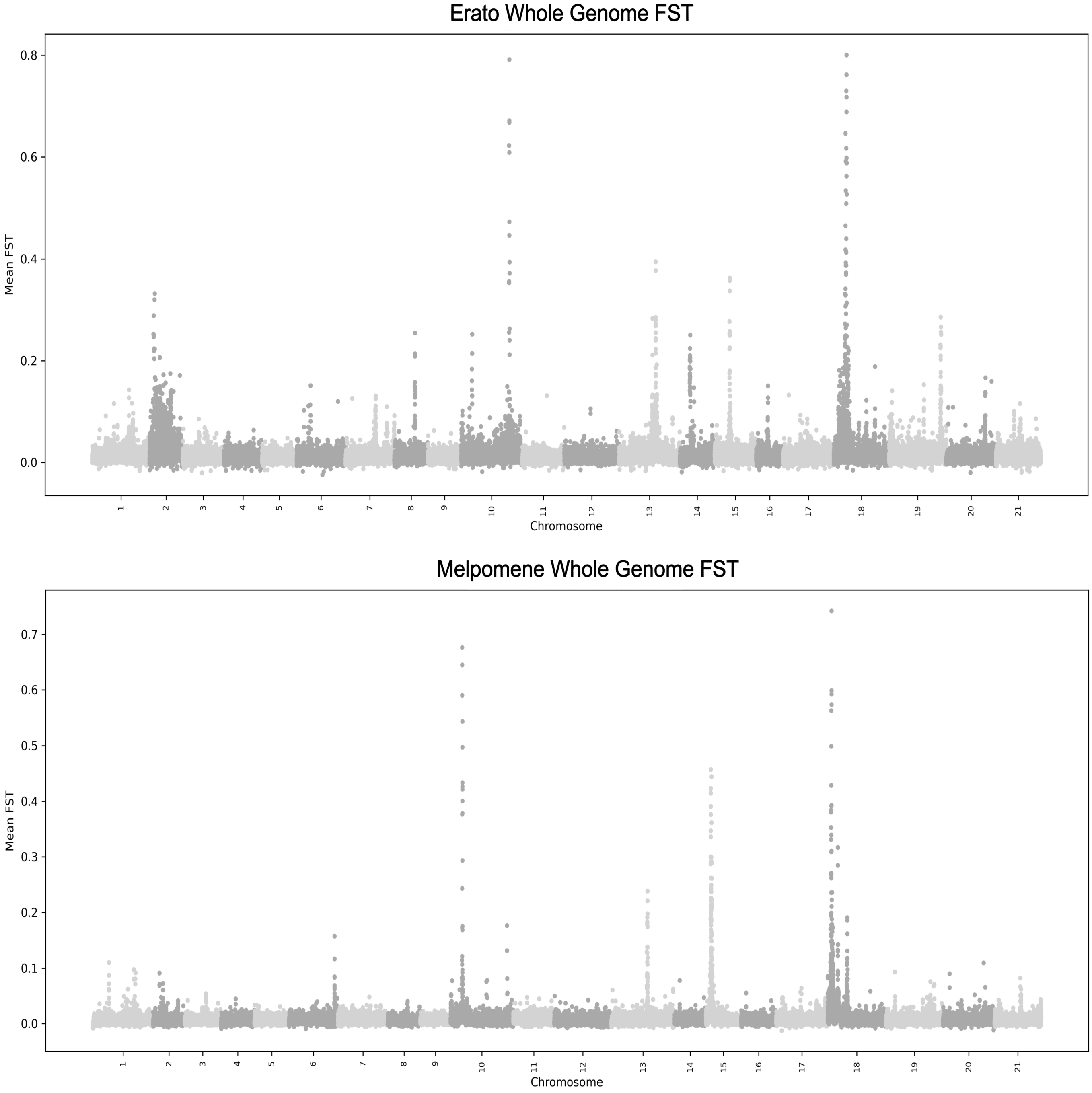


**Supplementary Figure 6. Whole-genome FST landscapes of *Heliconius erato* and *H. melpomene*.** Manhattan plots of genome-wide FST are shown for *H. erato* (top) and *H. melpomene* (bottom). Elevated differentiation is observed at multiple regions throughout the genome. Many of the major peaks coincide with known color pattern genes or genomic regions associated with wing patterning.
